# Meta-analysis of MERS, SARS and COVID-19 *in vitro* Infection Datasets Reveals Common Patterns in Gene and Protein Expression

**DOI:** 10.1101/2020.09.25.313510

**Authors:** Axel Martinelli, Murodzhon Akhmedov, Ivo Kwee

## Abstract

Three lethal lower respiratory tract coronavirus epidemics have occurred over the past 20 years. This coincided with major developments in genome-wide gene and protein expression analysis, resulting in a wealth of datasets in the public domain. Seven such in *vitro* studies were selected for comparative bioinformatic analysis through the VirOmics Playground, a visualisation and exploration platform we developed. Despite the heterogeneous nature of the data sets, several commonalities could be observed across studies and species. Differences, on the other hand, reflected not only variations between species, but also other experimental variables, such as cell lines used for the experiments, infection protocols and viral strains. The datasets analysed here are available online through our platform (https://public.bigomics.ch/app/omicsplayground_viromics).

## Introduction

Common human Coronavirus species, of which four are currently known, are usually associated with infections of the upper respiratory tract and cold-like symptoms [1]. However, the emergence of species infecting the lower respiratory tract and causing lethal epidemics has been a recurring event for the past 20 years. Starting with the SARS epidemic in China in 2002, the MERS epidemic in the Middle East ten years later and finally to the current COVID-19 pandemic [2].

The causative agents are all single-stranded RNA viruses (MERS-CoV for MERS, SARS-CoV-1 for SARS and SARS-CoV-2 for COVID-19) belonging to the beta-coronavirus genus within the Coronaviridae family [3]. All three infections started as zoonotic diseases and while the animal reservoir for both SARS-CoV-1 and MERS-CoV have been discovered, there is still uncertainty concerning the animal host for SARS-CoV-2, although it is most closely related to a bat SARS-like coronavirus [4]. SARS-CoV-1 and SARS-CoV-2 are, as the name implies, also genetically closer to each other than to MERS-CoV. This is reflected in the receptors that are recognised by the viral spikes surface glycoproteins of each virus. Thus, while MERS-CoV binds to dipeptidyl peptidase 4 (DPP4) on the surface of host cells [5], both SARS-CoV-1 and SARS-CoV-2 use the Angiotensin-converting enzyme 2 (ACE2) as the main entry point into the cell [6, 7]. Binding to ACE2 is induced by the priming of the viral S protein via the human serine protease TMPRSS2 [7]. A study on mice showed that SARS-CoV-1 infections resulted in the reduction of ACE2 expression *in vivo* [8], which was postulated to be responsible for several of the pathologies induced by the infection.

The epidemiologies of the three diseases display different characteristics. While MERS was a highly lethal infection with a low reproductive rate, both SARS and COVID-19 displayed a similarly high reproductive rate (>3) but a lower fatality rate [3]. COVID-19 has by far the lowest mortality rate of the three and also the highest estimated reproductive rate, which may explain why it is the only one of the three viruses to cause a global pandemic.

The clinical manifestations caused by the three viruses are similar. They all affect the lungs and provoke extensive alveolar damage [2]. However, while patients affected with SARS or MERS almost always developed a fever and also presented a high number of cases with gastrointestinal symptoms, patients with COVID-19 rarely develop gastrointestinal symptoms. Furthermore, the majority of COVID-19 patients are either asymptomatic or present mild symptoms, a factor that has played a role in the spread of the disease [3]. Thrombosis and embolisms have been observed both in SARS and COVID-19 patients [9, 10] and may explain sudden deaths observed in recovering patients.

All three infections have been associated with a cytokine storm triggered by an excessively stimulated immune system [3], which contributes to the morbidity of the diseases in severe patients. This cytokine storm is characterised by the release of large quantities of pro-inflammatory cytokines (such as Interferon-alpha (IFN-alpha), IFN-gamma, Tumor Necrosis Factor alpha (TNF-alpha) and Interleukin 6 (IL-6)) and chemokines, which induce an immune responses that damages internal organs and can result in death [11]. Intriguingly, the upregulation of anti-inflammatory cytokines such as IL-4 and IL-10 has also been observed [12]. The use of cytokine inhibitors may be beneficial after the onset of the cytokine storm. In particular, tocilizumab, an anti-IL-6 receptor monoclonal antibody, has been tested in COVID-19 patients and showed promising results [13]. Conversely, IFN-beta expression is inhibited by SARS-CoV-1 infections [14], while a general type I interferon inhibition has been observed in SARS-CoV-2 infections [15]. Type I interferons have antiviral properties and may be actively repressed by SARS-CoV-1 and SARS-CoV-2 during infection to escape the host immune response. Indeed, a clinical trial with an IFN-beta aerosol indicated a reduction in the number of patients developing critical conditions [16].

Various antiviral and anti-inflammatory drugs have been tested for their efficacy against coronavirus infections. Among them, the most prominent ones are: lopinavir/ritonavir, remdesivir, chloroquine [3], baricitinib, fedratinib, and ruxolitinib [17, 18]. Trametinib and selumetinib, two ERK/MAPK signal pathway inhibitors, have also shown significant efficacy in *vitro* against MERS-CoV infections [19].

These epidemics coincided with the development and rise of large scale gene expression analysis technologies (microarray and later on RNA-seq), with various groups embracing these technologies to generate a plethora of data sets that are currently publicly available.

For this review, we have selected seven public datasets (six transcriptomes and a proteome) based on cell lines infected with MERS-CoV, SARS-CoV-1 and SARS-CoV-2. While not aiming at providing a comprehensive meta-analysis, we wanted to highlight major similarities and differences both across species and across studies on the same species at the gene expression or proteomic level.

## Materials and methods

We selected a total of seven studies: six consisted of transcriptomic data (four microarrays, two RNA-seq) and one consisted of proteomic data. Of these studies three were conducted on MERS-CoV, one on SARS-CoV-1, one on SARS-CoV-2 and one on both SARS-CoV-1 and SARS-CoV-2. Datasets were obtained from the Gene Expression Omnibus (GEO) repository, except for the proteomic dataset, which was obtained from the Proteomics Identification Database (PRIDE). Details on each dataset are provided in Table 1. Where available we used the read or LC-MS/MS spectral counts produced by the authors, otherwise we generated read counts from the raw fastq files with the Salmon software [20].

**Table 1.** Details of the datasets used in the analysis.

| GEO/PXD Code | Data Type | Virus | Cell Line | Published |
| --- | --- | --- | --- | --- |
| GSE45042 | Microarray | MERS-CoV | Calu-3 | [21, 22] |
| GSE65574 | Microarray | MERS-CoV | Calu-3 | [23] |
| GSE86529 | Microarray | MERS-CoV | HF | NA |
| GSE37827 | Microarray | SARS-CoV-1 | Calu-3 | NA |
| GSE147507 | RNA-seq | SARS-CoV-2 | NHBE, A459 | [24] |
| GSE148729 | RNA-seq | SARS-CoV-1/2 | Caco-2, Calu-3, H1299 | NA |
| PXD017710 | Proteome | SARS-CoV-2 | Caco-2 | [25] |

Table notes. A459: adenocarcinomic human alveolar basal epithelial cells; Caco-2: Caucasian colon adenocarcinoma; Calu-3: human non-small-cell lung cancer cell; H1299: human non-small cell lung carcinoma cell ; HF: human fibroblasts; NHBE: normal bronchial epithelial cells.

For the analysis presented here, only measurements taken 24h or longer post-infection were considered. For the GSE65574 dataset, only the infection with the wildtype MERS virus was considered. In total, the analysed samples consisted of five samples infected with MERS-CoV, eleven samples infected with SARS-CoV-1 and seven samples (including the proteome dataset) infected with SARS-CoV-2. The filtered read counts were then analysed with VirOmics Playground, a cloud-based open access bioinformatics interface based on a platform that we developed for the analysis and visualisation of transcriptomic and proteomic data [26].

Unless otherwise stated, significant differential expression (q<0.05) was calculated using the intersection between three algorithms, namely edgeR, DEseq2 and limma [27–29], and with a minimum logFC of 0.5. The same approach was also used when identifying significantly altered KEGG pathways, when performing a drug connectivity analysis or when performing a word cloud analysis, with the three algorithms being camera, GSVA and fGSEA [30–32].

Discovery of pathways with altered expression for the proteomic data was performed via an enrichment analysis of keywords in reference datasets (displayed as a world cloud), since the KEGG pathway analysis with combined algorithms did not yield significant results. More information on the methods used can be found both in Akhmedov et al [26] and on the online documentation (https://omicsplayground.readthedocs.io/en/latest/).

The L1000 Connectivity Map database [33] was accessed via the platform to identify common inhibitory drug mode of actions against all three viral species, as described in Akhmedov et al [26].

Venn diagrams of the KEGG pathways comparisons were generated with an online tool (http://bioinformatics.psb.ugent.be/webtools/Venn/).

## Results

### Differentially Expressed Genes

The numbers of differentially expressed genes (or proteins) varied greatly across studies. While the numbers were fairly similar across the three MERS datasets, great variations were observed in both the SARS-CoV-1 and SARS-CoV-2 datasets (Table 2). Thus, for example, the number of differentially expressed genes in the SARS-CoV-2 datasets (excluding the H1299 cell line that yielded no dysregulated genes) ranged from 55 for the GSE147507 A459 cell line to 4425 for the GSE148729 Calu-3 cell line. This could be a reflection of the different cell lines, infection protocols and possibly viral strains used in the studies. It was also observed that the H1299 cell line (potentially a typical host cell for SARS viruses) did not show any evidence of gene expression alterations. This could be due to a failure in the infection protocol.

**Table 2.** Dysregulated genes (or proteins) in the different groups from the selected studies.

| **SAMPLE** | **UP** | **DOWN** |
| --- | --- | --- |
| COVID-19 Proteome 24h | 177 | 261 |
| GSE45042 MERS 24h | 4264 | 4450 |
| GSE65574 icMERS 24h | 3003 | 2905 |
| GSE86529 MERS 24h | 4045 | 4367 |
| GSE86529 MERS 36h | 3330 | 3189 |
| GSE86529 MERS 48h | 3347 | 3290 |
| GSE37827 icSARS 24h | 51 | 21 |
| GSE37827 icSARS 30h | 210 | 31 |
| GSE37827 icSARS 36h | 475 | 95 |
| GSE37827 icSARS 48h | 1853 | 1808 |
| GSE37827 icSARS 54h | 1794 | 2031 |
| GSE37827 icSARS 60h | 1910 | 2256 |
| GSE37827 icSARS 72h | 1677 | 1793 |
| GSE147507 COVID-19 A549 24h | 53 | 2 |
| GSE147507 COVID-19 NHBE 24h | 135 | 41 |
| GSE148729 SARS H1299 24h | 0 | 0 |
| GSE148729 COVID-19 H1299 24h | 0 | 0 |
| GSE148729 SARS H1299 36h | 0 | 0 |
| GSE148729 COVID-19 H1299 36h | 0 | 0 |
| GSE148729 SARS Calu-3 | 879 | 956 |
| GSE148729 COVID-19 Calu-3 | 2141 | 2294 |
| GSE148729 SARS Caco-2 | 1186 | 1086 |
| GSE148729 COVID-19 Caco-2 | 244 | 101 |

Table notes. The numbers were based on the intersection of three algorithms with $\log_{2}FC > 0.5$ and $q < 0.05$ , except for the proteome dataset where a single algorithm (DEseq2) and a $\log_{2}FC$ of 0.2 were selected.

In the only experiment where the same cell lines were infected with SARS-CoV-1 and SARS-CoV-2, there were striking differences in the amount of differentially expressed genes. Thus, in Caco-2 cells, infections with SARS-CoV-1 generated almost 6x more dysregulated genes compared to an infection with SARS-CoV-2. Meanwhile, the opposite was true in Calu-3 cells, where a SARS-CoV-2 infection generated more than 2X differentially expressed genes compared to a SARS-CoV-1 infection. Given the genetic similarity between the two viruses, such differences are rather unexpected.

When looking at the top 10 up- and down-regulated genes in each sample split by virus (excluding samples consisting of H1299 infected cells), no shared downregulated genes could be observed across virus species. However, MERS-CoV infected cells all shared the same most upregulated gene (SSX2), independently of time of collection or cell type. Furthermore, gene SLC22A23 was identified among the top 10 upregulated genes in all the samples collected 24h post infection. Gene FBXL8 (F-box and leucine rich repeat protein 8) was in the top 10 upregulated genes in all samples from datasets GSE65574 and GSE86529 and was also significantly upregulated in dataset GSE45042. Gene HSPA6 was in the top 10 upregulated genes of all the GSE86529 samples and was also significantly upregulated in the samples from the other two datasets.

For SARS-CoV-1 infected Calu-3 cells, IL-6 was among the top 10 upregulated genes across various time points in both available studies. For SARS-CoV-2 infected cells, no shared upregulated genes could be found. Considering that all available samples were from a different cell line each may explain the lack of commonality.

### Viral spike protein cellular targets and priming protein

There was no consistent up- or down-regulation of the DPP4 gene (the receptor for the MERS-CoV S protein) across the seven studies (Fig 1-3 A-C). In MERS-infected samples, the gene was significantly upregulated in study GSE45042 and non-significantly in study GSE65574 (Fig 1A and 1B), but downregulated in study GSE86529 (Fig 1C). One SARS-CoV study (GSE148729) conversely indicated down-regulation in Calu-3 cells and, in the case of SARS-CoV-1, Caco-2 cells (Fig 2B and 3A). There was no statistically significant change in expression in other SARS-CoV infected samples.

**Fig 1.**
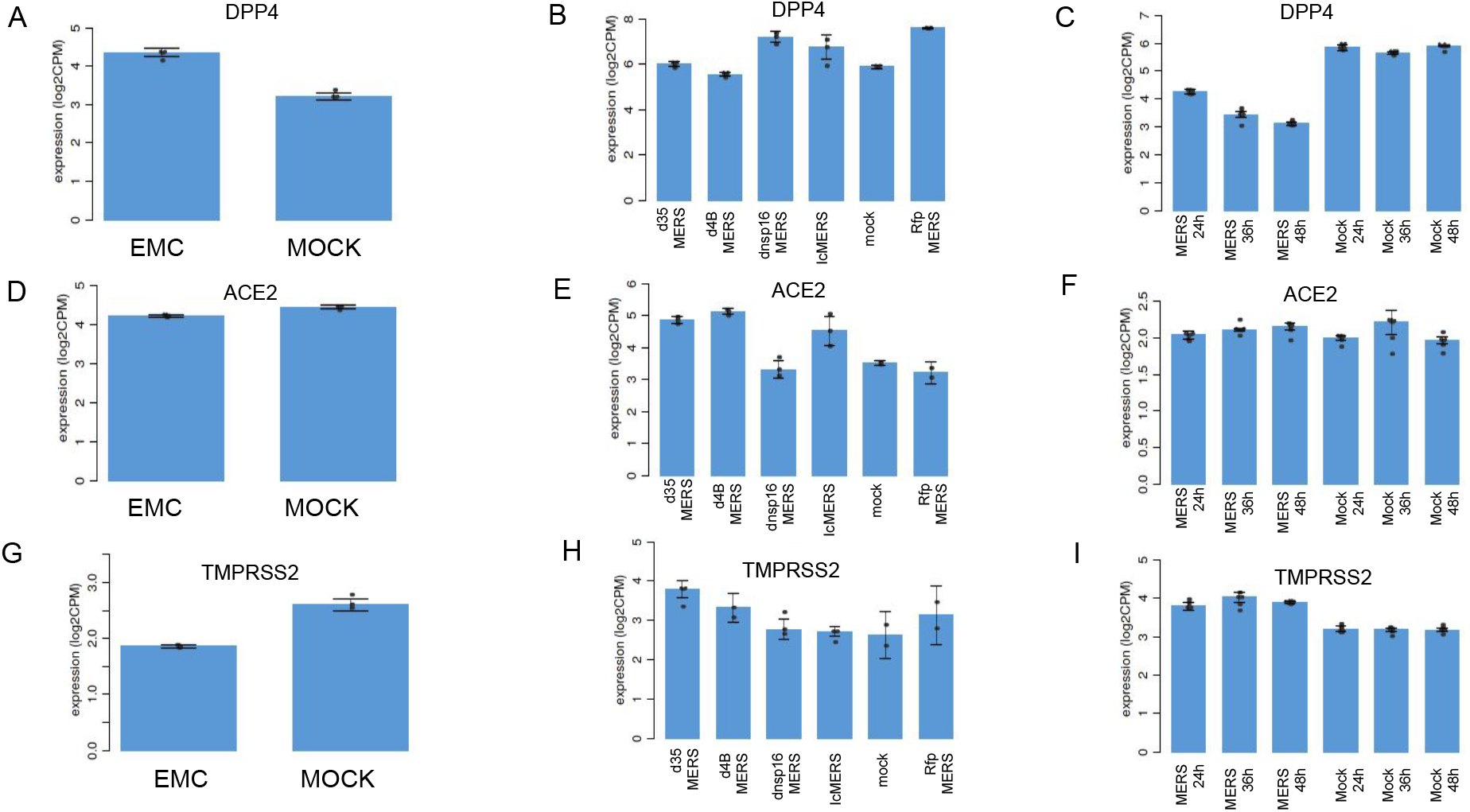
Expression levels in MERS-CoV infected cells. Expression levels for genes DPP4, ACE2 and TMPRSS2 per sample groups expressed in logCPM (count per million) from the three MERS-CoV infected datasets. A,D,G: GSE45042; B,E,H: GSE65574; C,F,I: GSE86529.

**Fig 2.**
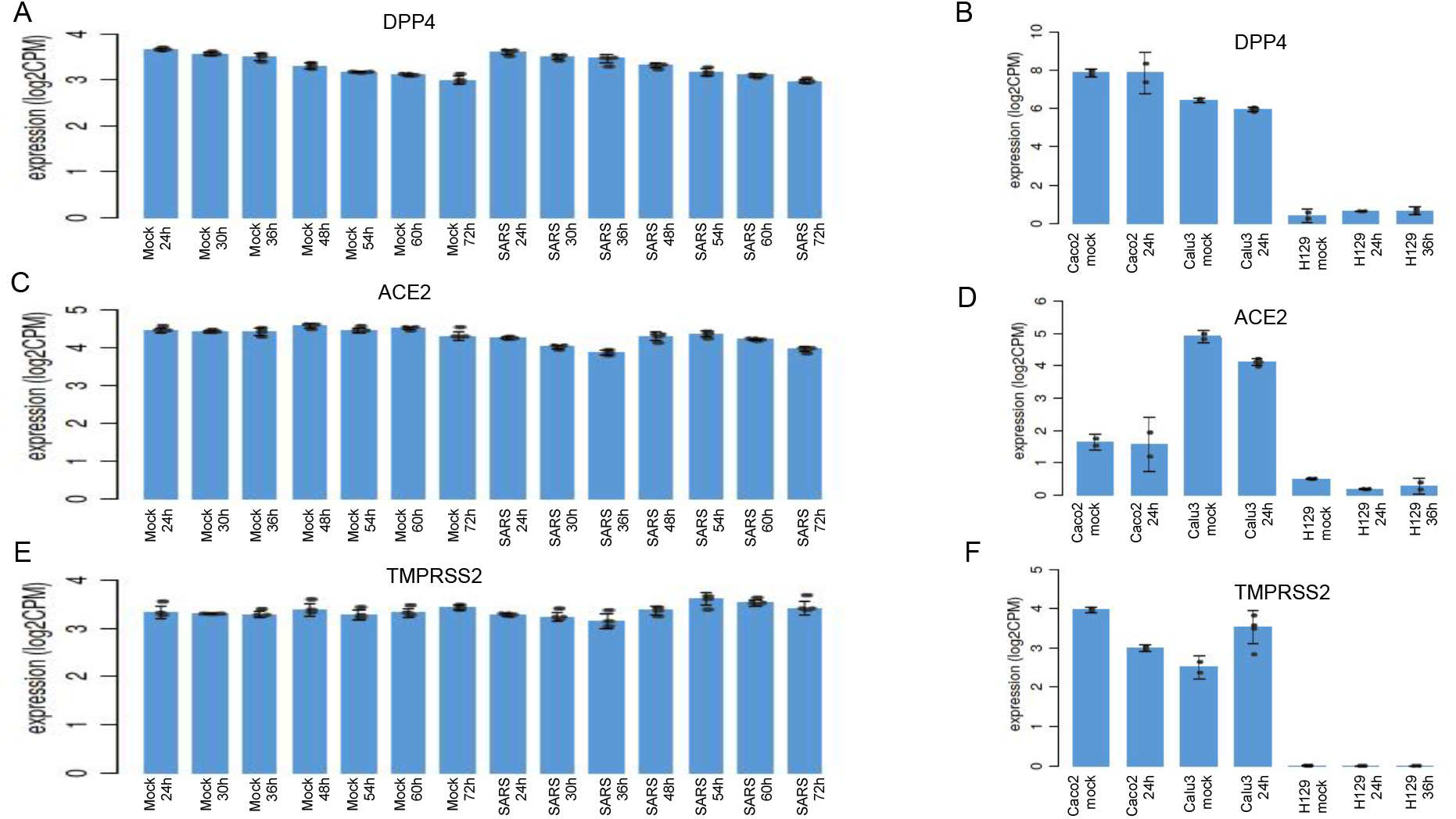
Expression levels in SARS-CoV-1 infected cells. Expression levels for genes DPP4, ACE2 and TMPRSS2 per sample groups expressed in logCPM (count per million) from two SARS-CoV-1 infected datasets. A,C,E: GSE37827; B,D,F: GSE148729.

**Fig 3.**
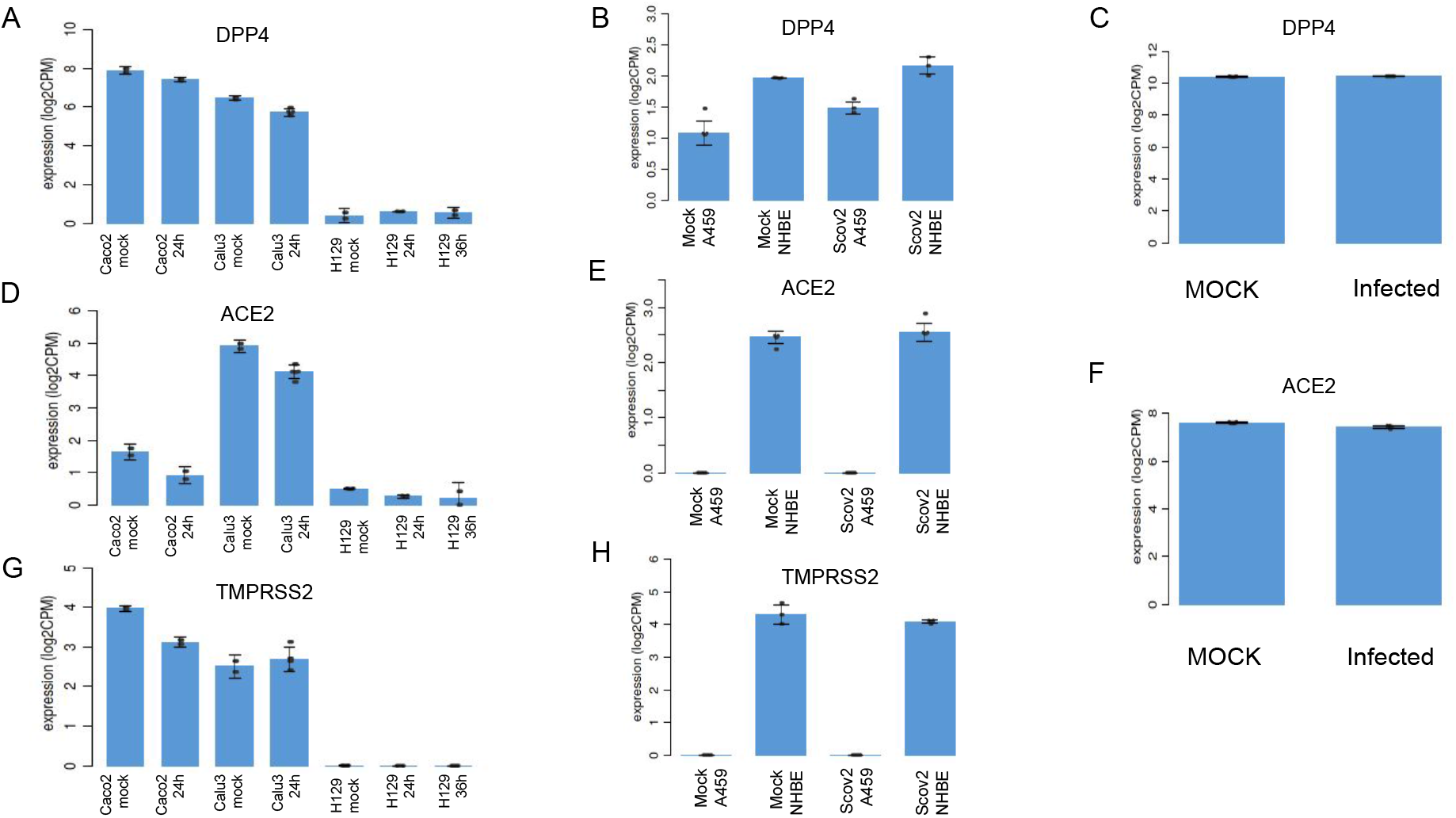
Expression levels in SARS-CoV-2 infected cells. Expression levels for genes DPP4, ACE2 and TMPRSS2 per sample groups expressed in logCPM (count per million) from three SARS-CoV-2 infected datasets. A,D,G: GSE147507; B,E,H: GSE148729.; C,F: proteome.

The expression of the ACE2 gene, whose product serves as a receptor for the SARS-CoV-1/2 S protein, was unaltered in MERS infected cell lines (Fig 1D-F). The picture was more mixed in SARS-CoV-1/2 infected cell lines. There was no evidence of gene expression reduction in Calu-3 cells infected with SARS-CoV-1 (Fig 2C) or NHBE and A459 cells infected with SARS-CoV-2 (Fig. 3E). Even at the proteome level, no difference in expression was detected (Fig 3F). Only in one study, infection with both SARS-CoV species induced a statistically significant reduction in gene expression, but only in one of the three cell lines that were infected, namely Calu-3 (Fig 2D and 3B).

Finally, TMPRSS2 gene expression provided very mixed results. It was observed to be either down-regulated (Fig 1G), up-regulated (Fig 1I) or unaltered (Fig 1H) in MERS-infected cell lines. Expression was down-regulated in Calu-3 cells infected with either SARS-CoV-1 or SARS-CoV-2 (Fig 2F and 3G), but was otherwise unaffected in all other samples. No data was available from the proteome study.

### Immune Response Genes

Coronavirus infections have been associated with the increased expression of various cytokines. Fifteen cytokines selected based on the available literature was selected for analysis across any of the available samples collected at least 24h post infection, excluding the proteomic dataset as none of the cytokines were present there, and the H1299 cell lines, as there were no dysregulated genes present, for a total of 18 samples.

Of these cytokines, seven were found to be upregulated in at least one sample for each of the three species, namely: IL-1B, IL-6, IL-8, IL-12A, CXCL10 (C-X-C motif chemokine ligand 10), CCL2 (Chemokine C-C ligand 2) and TNF-alpha (Fig 4). IL-6, and to a lesser extent IL-8 and CCL2, were frequently upregulated across all three diseases (MERS, SARS and COVID-19). IL-7, on the other hand, was the only gene upregulated in SARS and COVID-19, but not MERS.

**Fig 4.**
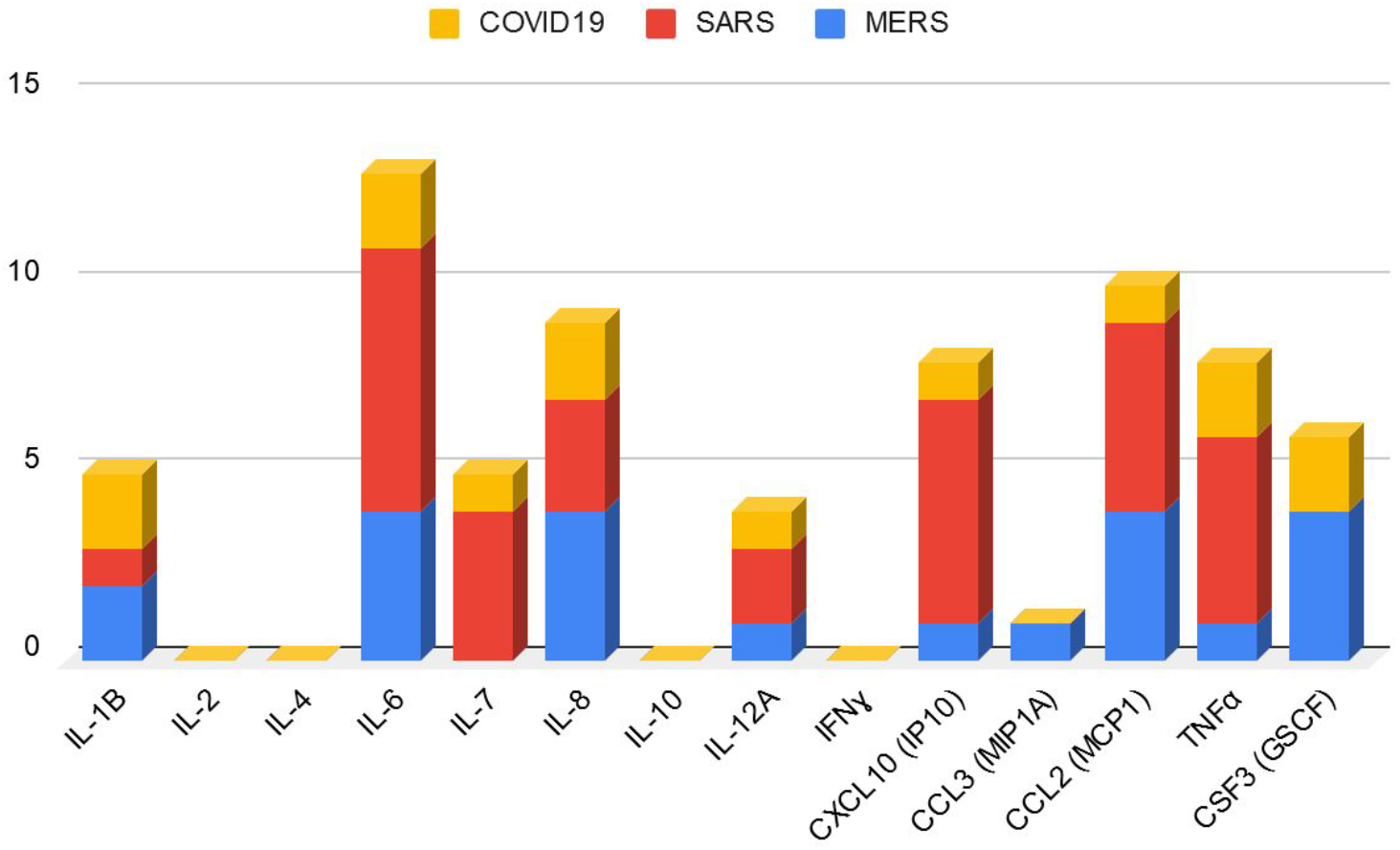
Upregulated cytokines. Instances of significantly upregulated cytokines in any sample collected 24h or more post infection, divided by disease.

IFN-beta and IFN-epsilon were the only two type I interferons that were measured in all three SARS and COVID-19 datasets (GSE14707, GSE37827 and GSE148729). There was a complete lack of statistically significant upregulation for IFN-epsilon following infection in any of the datasets at any available time point (Fig 5-7). While there was a lack of upregulation early on in the infection, after 24h IFN-beta expression did increase in Calu-3 cells in two of the datasets (Fig 5-6). In the only study where SARS-CoV-2 infections were compared against two other respiratory tract viruses, namely Influenza A virus (IAV) and Respiratory Syncytial Virus (RSV), the difference in type I interferon expression levels was striking.SARS-CoV-2 infections showed expression profile indistinguishable from control samples, in clear contrast with a general upregulation of the same interferons in the infections with IAV or RSV (Fig 7E-H). The picture was less clear in MERS-CoV infected samples, with some showing a slight upregulation and others no change in Type I interferon expression levels (data not shown).

**Fig 5.**
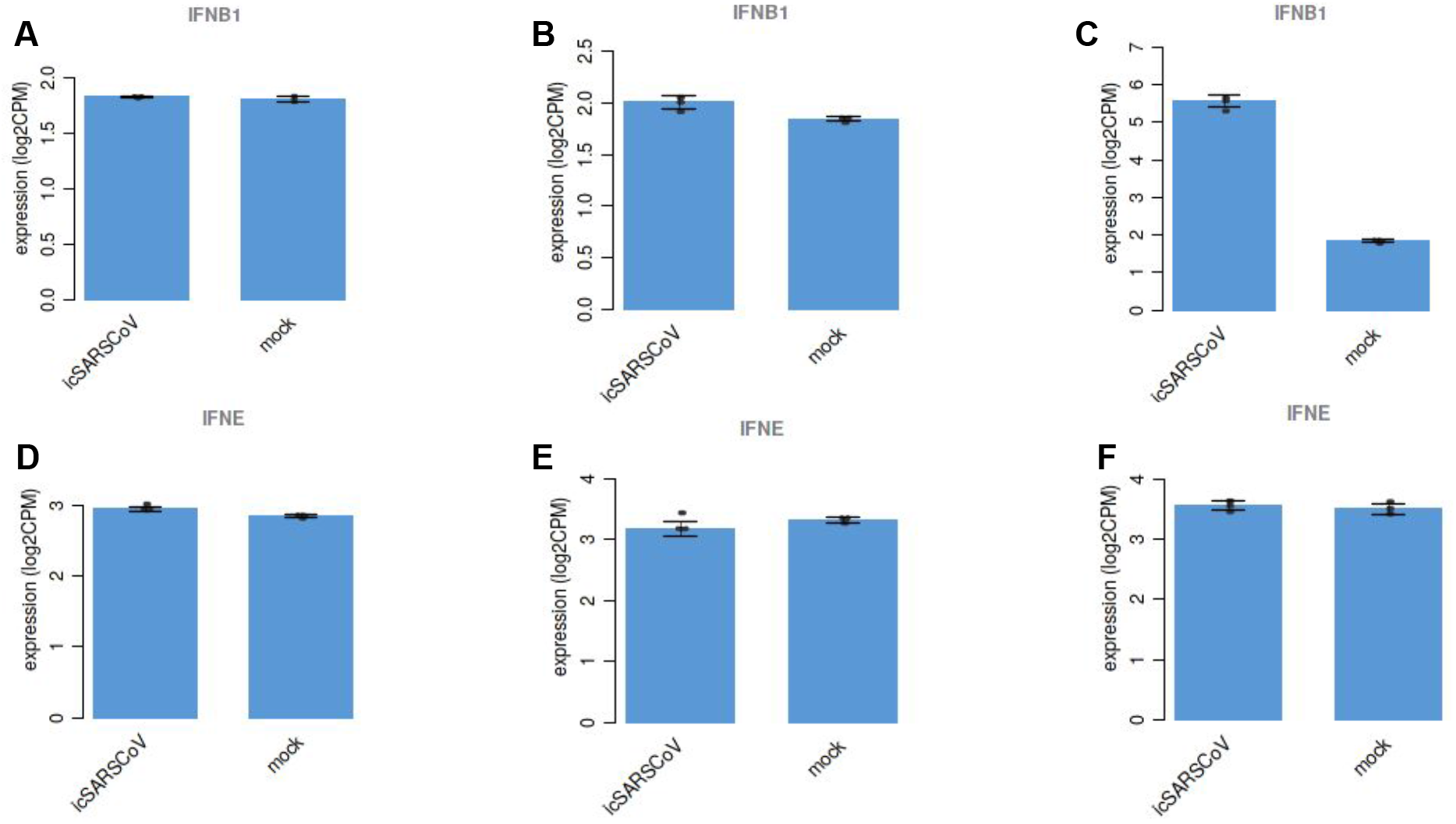
Type I interferon gene expression in samples infected with SARS-CoV-1 from study GSE37827. A: IFN-beta expression in Calu-3 cells 12h post-infection; B: IFN-beta expression in Calu-3 cells 24h post-infection; C: IFN-beta expression in Calu-3 cells 48h post-infection; D: IFN-epsilon expression in Calu-3 cells 12h post-infection; E: IFN-epsilon expression in Calu-3 cells 24h post-infection; F: IFN-epsilon expression in Calu-3 cells 48h post-infection.

**Fig 6.**
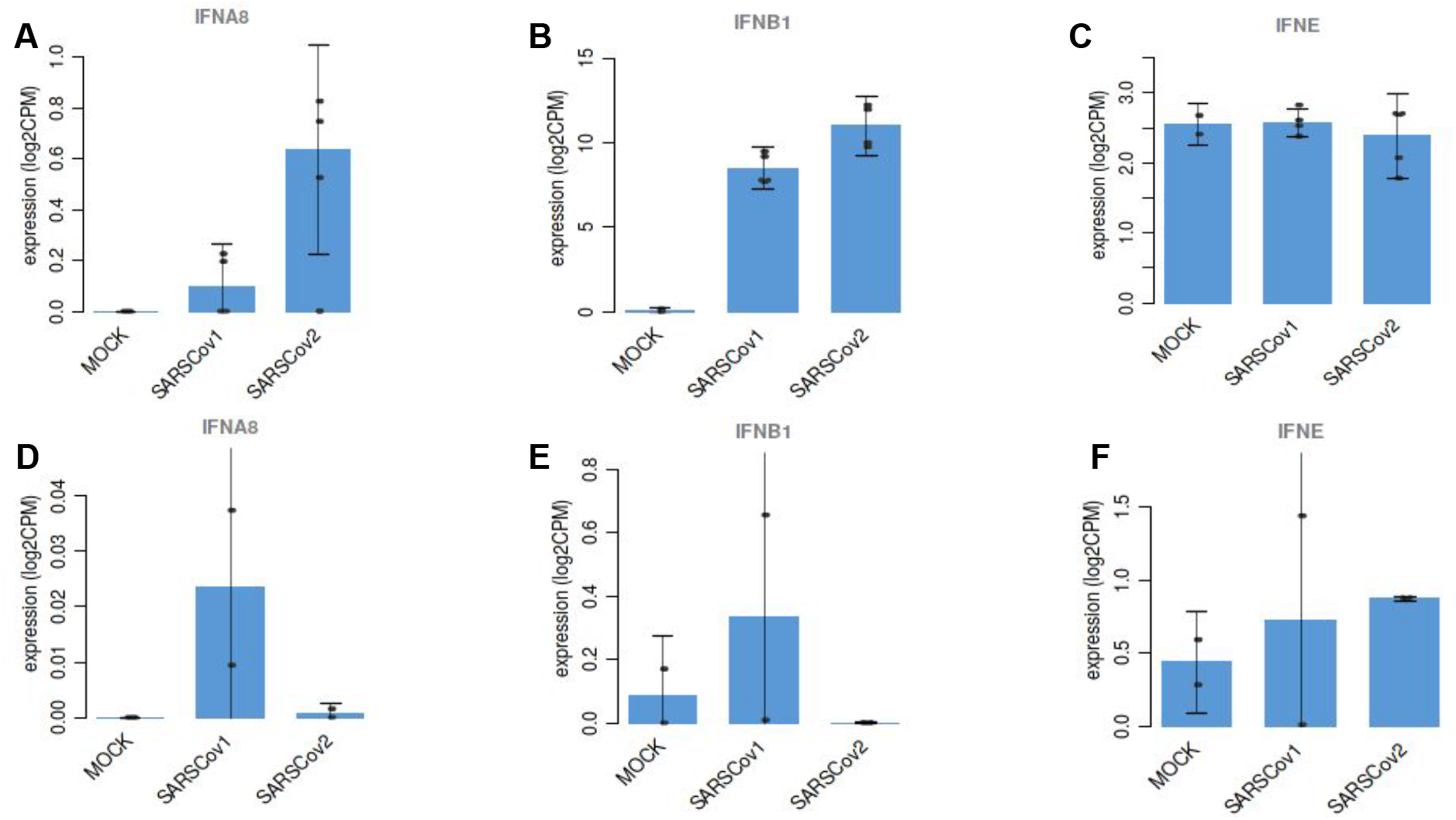
Type I interferon gene expression in samples infected with SARS-CoV-1 and SARS-CoV-2 from study GSE148729. A: IFN-alpha-8 expression in Calu-3 cells; B: IFN-beta expression in Calu-3 cells; C: IFN-epsilon expression in Calu-3 cells; D: IFN-alpha-8 expression in Caco-2 cells; E: IFN-beta expression in Caco-2 cells; F: IFN-epsilon expression in Caco-2 cells.

**Fig 7.**
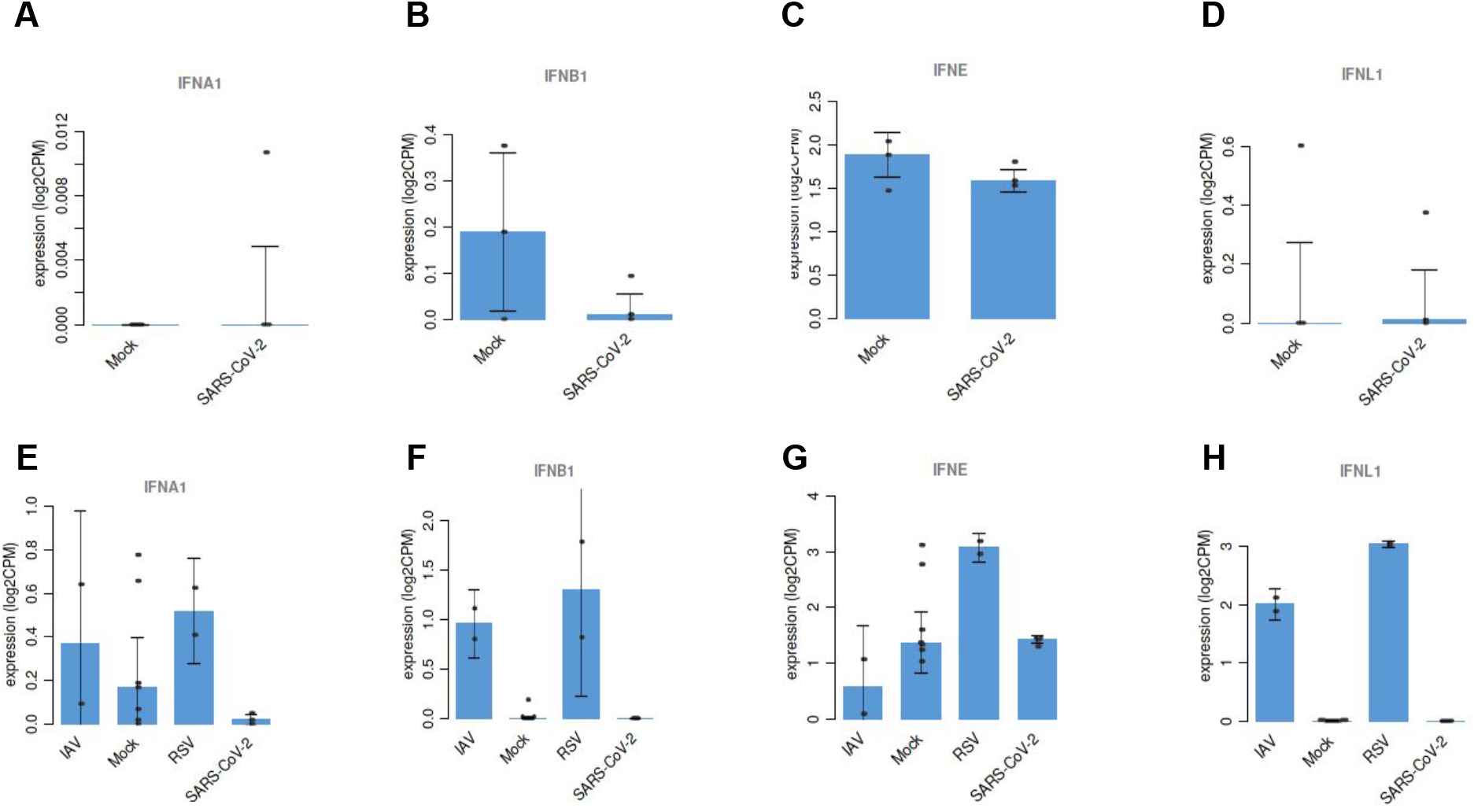
Type I interferon gene expression in samples infected with SARS-CoV-2, RSV and IAV from study GSE147507. A: IFN-alpha-1 expression in NHBE cells; B: IFN-beta expression in NHBE cells; C: IFN-epsilon expression in NHBE cells; D: IFN-lambda-1 expression in NHBE cells; E: IFN-alpha-1 expression in A459 cells; F: IFN-beta expression in A459 cells; G: IFN-epsilon expression in A459 cells; H: IFN-lambda-1 expression in A459 cells.

### Altered Pathway Analysis

For the six transcriptome studies, a KEGG pathway analysis was performed to identify pathways with significantly (q<0.05) altered expression levels in all samples collected at least 24h post infection. Samples composed of infected H1299 cell lines were excluded, due to the lack of differentially expressed genes. As the KEGG pathway analysis yielded no significant results for the proteome dataset, its protein expression pattern was compared against a collection of more than 50,000 genesets contained within the platform [26] and a word cloud plot based on the results was generated.

The word cloud plot (Fig 8) contained terms that have been described in the published article based on the same dataset [25]. Thus, the term “cholesterol” appears with a negative normalised enrichment score (NES), indicating a downregulation of associated genes. Conversely, genes in keywords associated with carbon metabolism (glycolysis, glucose) were upregulated. A slight upregulation of the spliceosome KEGG pathway could be observed, although it was not statistically significant (data not shown).

**Fig 8.**
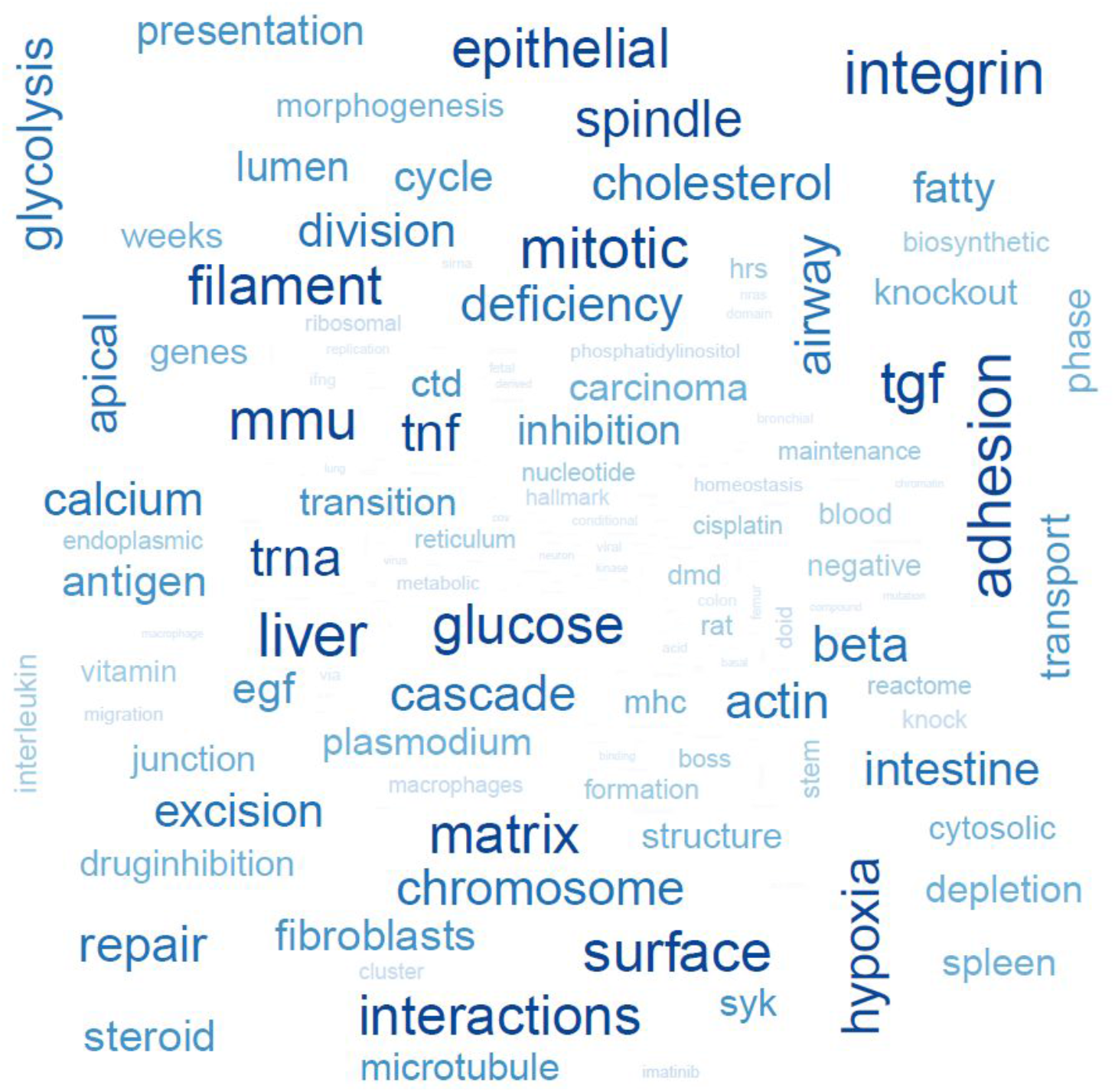
Word cloud plot of the most common keywords in the proteome dataset analysis. Word cloud of the most recurring keywords in the top ranked gene sets used to compare the differentially expressed genes appearing 24h post infection in the proteome study [25].

Comparing the various samples across studies is not obvious, because of the different conditions (viral species, time of collections, cell lines and infection protocols). The results of the various datasets were thus combined in two ways. First of all, only the samples collected 24h post infection were considered. The resulting differentially expressed pathways (Table S1-S3) were split into up- and down-regulated groups and combined by species and intersections between the three species displayed as Venn diagrams. Secondly, only experiments conducted on Calu-3 cells (the most common cell used in the studies selected here) were chosen, but including any samples collected between 24h and 72h post infection. The up- and down-regulated KEGG pathways (Table S4-S6) were again combined by species, and Venn diagrams were produced as indicated above.

The 24h samples intersection of KEGG pathways yielded three upregulated pathways and six downregulated that were shared by all three species (Fig 9-10), while two and eight were shared between SARS-CoV-1 and SARS-CoV-2 infected samples respectively (Fig 9-10). The upregulated pathways shared by all three species included terms related to immunity and ribosome formation, while downregulated pathways included the “Valine, Leucine and Isoleucine degradation” pathway, “pyruvate metabolism” and the “peroxisome” pathway (Table 3). The “complement and coagulation” pathway was among the upregulated pathways in common between SARS-CoV-1 and SARS-CoV-2 infected cells, while shared downregulated pathways included the “glycolysis and gluconeogenesis” pathway (Table 4).

**Fig 9.**
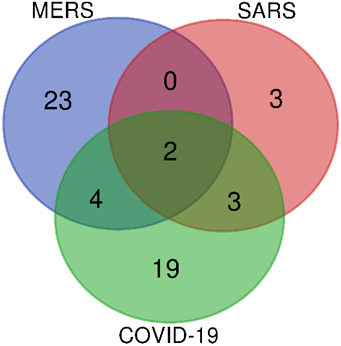
Venn diagram of upregulated pathways 24h post infection. Venn diagram representation of the intersection between upregulated pathways for infected samples collected 24h after infections.

**Fig 10.**
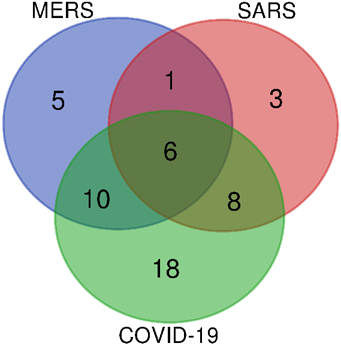
Venn diagram of downregulated pathways 24h post infection. Venn diagram representation of the intersection between downregulated pathways for infected samples collected 24h after infections.

**Table 3.** KEGG pathways shared between all three species in samples collected 24h post infections.

| KEGG Pathway | UP/DOWN Reg. |
| --- | --- |
| KEGG CYTOKINE CYTOKINE RECEPTOR INTERACTION | UP |
| PATHWAY:TNF signaling pathway Homo sapiens hsa04668 | UP |
| PATHWAY:Ribosome biogenesis in eukaryotes Homo sapiens hsa03008 | UP |
| C2:KEGG PROPANOATE METABOLISM | DOWN |
| C2:KEGG OXIDATIVE PHOSPHORYLATION | DOWN |
| C2:KEGG PARKINSONS DISEASE | DOWN |
| C2:KEGG VALINE LEUCINE AND ISOLEUCINE DEGRADATION | DOWN |
| C2:KEGG PYRUVATE METABOLISM | DOWN |
| C2:KEGG PEROXISOME | DOWN |

**Table 4.** KEGG pathways shared between SARS-CoV-1 and SARS-CoV-2 infected cells in samples collected 24h post infections.

| KEGG Pathway | UP/DOWN Reg. |
| --- | --- |
| PATHWAY:Osteoclast differentiation Homo sapiens hsa04380 | UP |
| C2:KEGG COMPLEMENT AND COAGULATION CASCADES | UP |
| C2:KEGG ASCORBATE AND ALDARATE METABOLISM | DOWN |
| C2:KEGG GLYCINE SERINE AND THREONINE METABOLISM | DOWN |
| C2:KEGG DNA REPLICATION | DOWN |
| C2:KEGG PENTOSE AND GLUCURONATE INTERCONVERSIONS | DOWN |
| C2:KEGG GLYCOLYSIS GLUCONEOGENESIS | DOWN |
| C2:KEGG METABOLISM OF XENOBIOTICS BY CYTOCHROME P450 | DOWN |
| C2:KEGG DRUG METABOLISM CYTOCHROME P450 | DOWN |
| C2:KEGG OOCYTE MEIOSIS | DOWN |

The intersection analysis based on Calu-3 cells, yielded 1 upregulated and 6 downregulated KEGG pathways shared between all viral species and 14 and 22 respectively shared between cells infected with SARS-CoV-1 and SARS-CoV-2 (Fig 11-12). The list of pathways shared by all three species included the “cytokine cytokine receptor” pathway, as well as pathways related to carbohydrate metabolism and protection from oxidation (Table 5). When comparing SARS-CoV-1 and SARS-CoV-2 infected Calu-3 cells, shared upregulated pathways included different immune-related gene sets, while shared carbohydrate metabolism-related pathways and butanoate metabolism were present among the downregulated pathways (Table 6).

**Fig 11.**
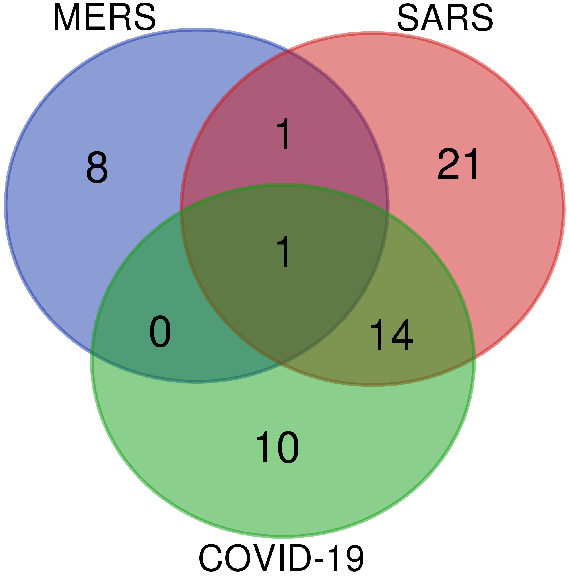
Venn diagram of upregulated pathways in infected Calu-3 cells 24h post infection. Venn diagram representation of the intersection between upregulated pathways for infected samples collected from Calu-3 infected cells.

**Fig 12.**
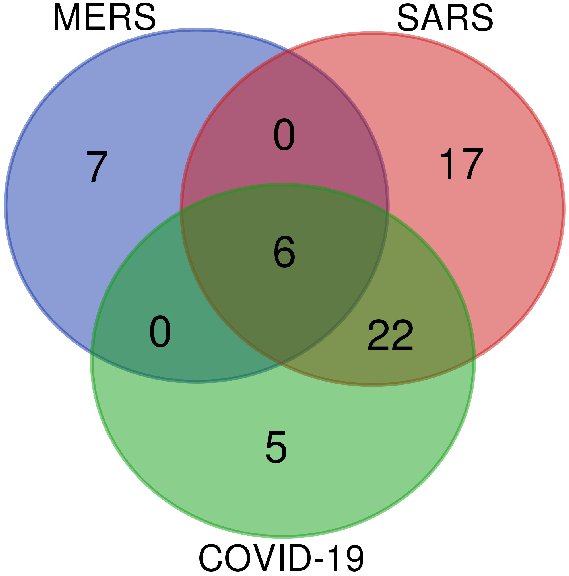
Venn diagram of downregulated pathways in infected Calu-3 cells 24h post infection. Venn diagram representation of the intersection between downregulated pathways for infected samples collected from Calu-3 infected cells.

**Table 5.** KEGG pathways shared between all three species in samples collected from Calu-3 infected cells.

| KEGG Pathway | UP/DOWN Reg. |
| --- | --- |
| KEGG CYTOKINE CYTOKINE RECEPTOR INTERACTION | UP |
| C2:KEGG FATTY ACID METABOLISM | DOWN |
| C2:KEGG LYSOSOME | DOWN |
| C2:KEGG CITRATE CYCLE TCA CYCLE | DOWN |
| C2:KEGG VALINE LEUCINE AND ISOLEUCINE DEGRADATION | DOWN |
| C2:KEGG PYRUVATE METABOLISM | DOWN |
| C2:KEGG GLUTATHIONE METABOLISM | DOWN |

**Table 6.** KEGG pathways shared between SARS-CoV-1 and SARS-CoV-2 infected Calu-3 cells.

| KEGG Pathway | UP/DOWN Reg. |
| --- | --- |
| KC2:KEGG RIG I LIKE RECEPTOR SIGNALING PATHWAY | UP |
| C2:KEGG GRAFT VERSUS HOST DISEASE | UP |
| PATHWAY:Herpes simplex infection Homo sapiens hsa05168 | UP |
| C2:KEGG TYPE I DIABETES MELLITUS | UP |
| C2:KEGG CYTOSOLIC DNA SENSING PATHWAY | UP |
| PATHWAY:TNF signaling pathway Homo sapiens hsa0466 | UP |
| PATHWAY:Influenza A Homo sapiens hsa05164 | UP |
| C2:KEGG JAK STAT SIGNALING PATHWAY | UP |
| C2:KEGG ANTIGEN PROCESSING AND PRESENTATION | UP |
| C2:KEGG NOD LIKE RECEPTOR SIGNALING PATHWAY | UP |
| PATHWAY:Staphylococcus aureus infection Homo sapiens hsa05150 | UP |
| C2:KEGG ALLOGRAFT REJECTION | UP |
| C2:KEGG TOLL LIKE RECEPTOR SIGNALING PATHWAY | UP |
| PATHWAY:Inflammatory bowel disease (IBD) Homo sapiens hsa05321 | UP |
| C2:KEGG PORPHYRIN AND CHLOROPHYLL METABOLISM | DOWN |
| C2:KEGG ASCORBATE AND ALDARATE METABOLISM | DOWN |
| C2:KEGG AMINO SUGAR AND NUCLEOTIDE SUGAR METABOLISM | DOWN |
| C2:KEGG PROPANOATE METABOLISM | DOWN |
| C2:KEGG FRUCTOSE AND MANNOSE METABOLISM | DOWN |
| C2:KEGG PENTOSE PHOSPHATE PATHWAY | DOWN |
| C2:KEGG ONE CARBON POOL BY FOLATE | DOWN |
| C2:KEGG STARCH AND SUCROSE METABOLISM | DOWN |
| C2:KEGG DNA REPLICATION | DOWN |
| C2:KEGG PENTOSE AND GLUCURONATE INTERCONVERSIONS | DOWN |
| C2:KEGG N GLYCAN BIOSYNTHESIS | DOWN |
| C2:KEGG GLYCOLYSIS GLUCONEOGENESIS | DOWN |
| C2:KEGG PARKINSONS DISEASE | DOWN |
| C2:KEGG CARDIAC MUSCLE CONTRACTION | DOWN |
| C2:KEGG ALZHEIMERS DISEASE | DOWN |
| C2:KEGG OOCYTE MEIOSIS | DOWN |
| C2:KEGG PROXIMAL TUBULE BICARBONATE RECLAMATION | DOWN |
| C2:KEGG METABOLISM OF XENOBIOTICS BY CYTOCHROME P450 | DOWN |
| C2:KEGG DRUG METABOLISM CYTOCHROME P450 | DOWN |
| C2:KEGG PEROXISOME | DOWN |
| C2:KEGG BUTANOATE METABOLISM | DOWN |
| C2:KEGG RETINOL METABOLISM | DOWN |

### Drug Connectivity Map Analysis

A Drug Connectivity analysis was performed on a total of five MERS-CoV infected samples, 9 SARS-CoV-1 infected samples and five SARS-CoV-2 infected samples (including the proteome study). Drug gene sets that had statistically significant inversely correlated expression profiles to the experimental results were obtained and summarised by mechanisms of action through our platform. The top 10 most negatively correlated mechanisms were then combined by disease (MERS, SARS or COVID-19).

Samples infected by MERS-CoV displayed a strong negative correlation with cyclin dependant kinase (CDK), epidermal growth factor receptor (EGFR), dual specificity mitogen-activated protein kinase (MEK), mammalian target of rapamycin (mTOR), phosphatidylinositol-3 kinases (PI3K) and tyrosine-protein kinase src (src) inhibitors, with at least 4 out of 5 samples showing a negative correlation with such mechanisms of action (Fig 13).

**Fig 13.**
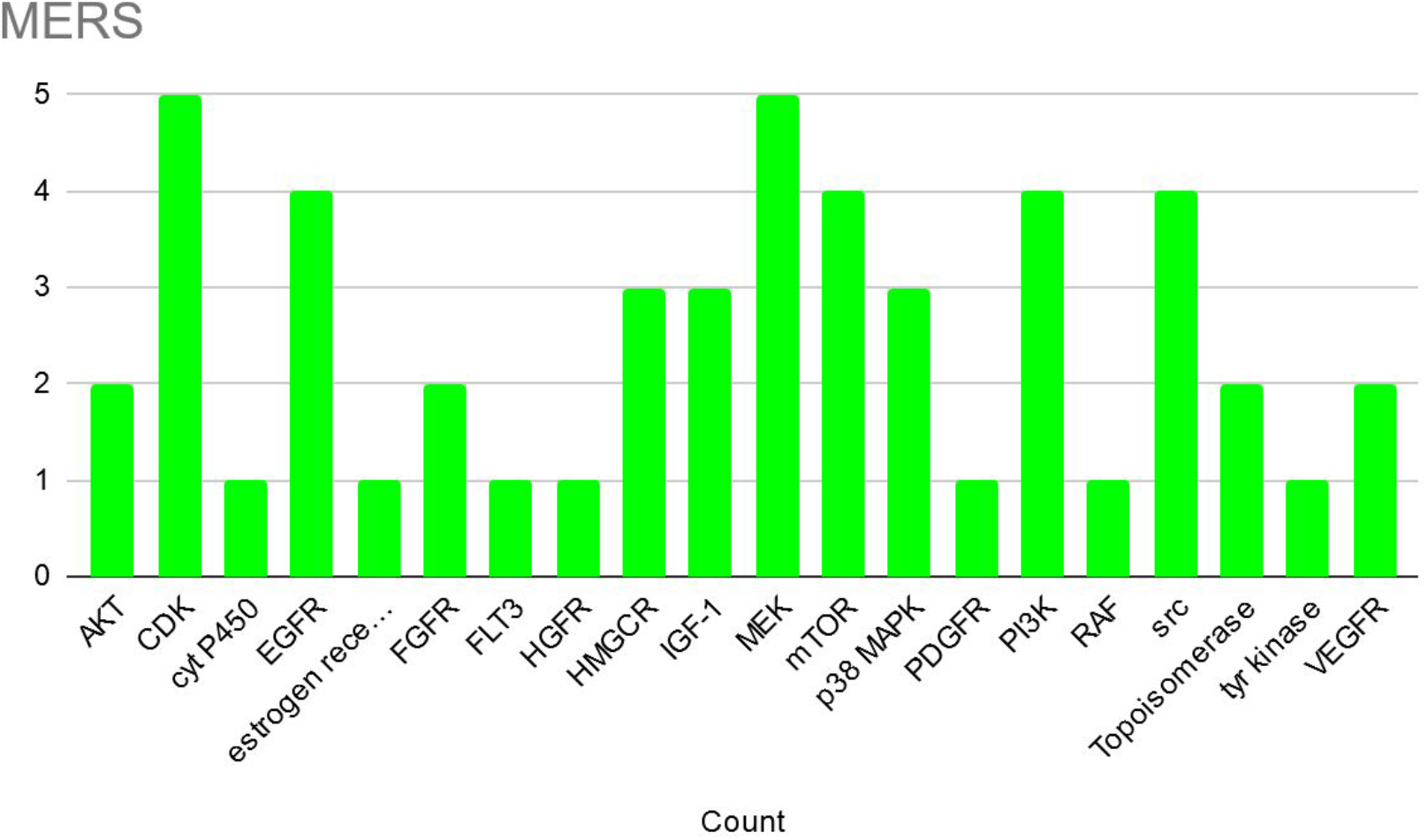
Anti-MERS-CoV inhibitory mechanisms. Cumulative counts of the most significant potential inhibitory mechanisms against MERS-CoV infections in five separate samples.

SARS-CoV-1 infected samples also showed a strong negative correlation (with detection among the top 10 inhibitory profiles in at least 7/9 samples) with CDK, EGFR, MEK and src inhibitors (Fig 14). Additionally, p38 MAPK inhibitors were also prominently represented.

**Fig 14.**
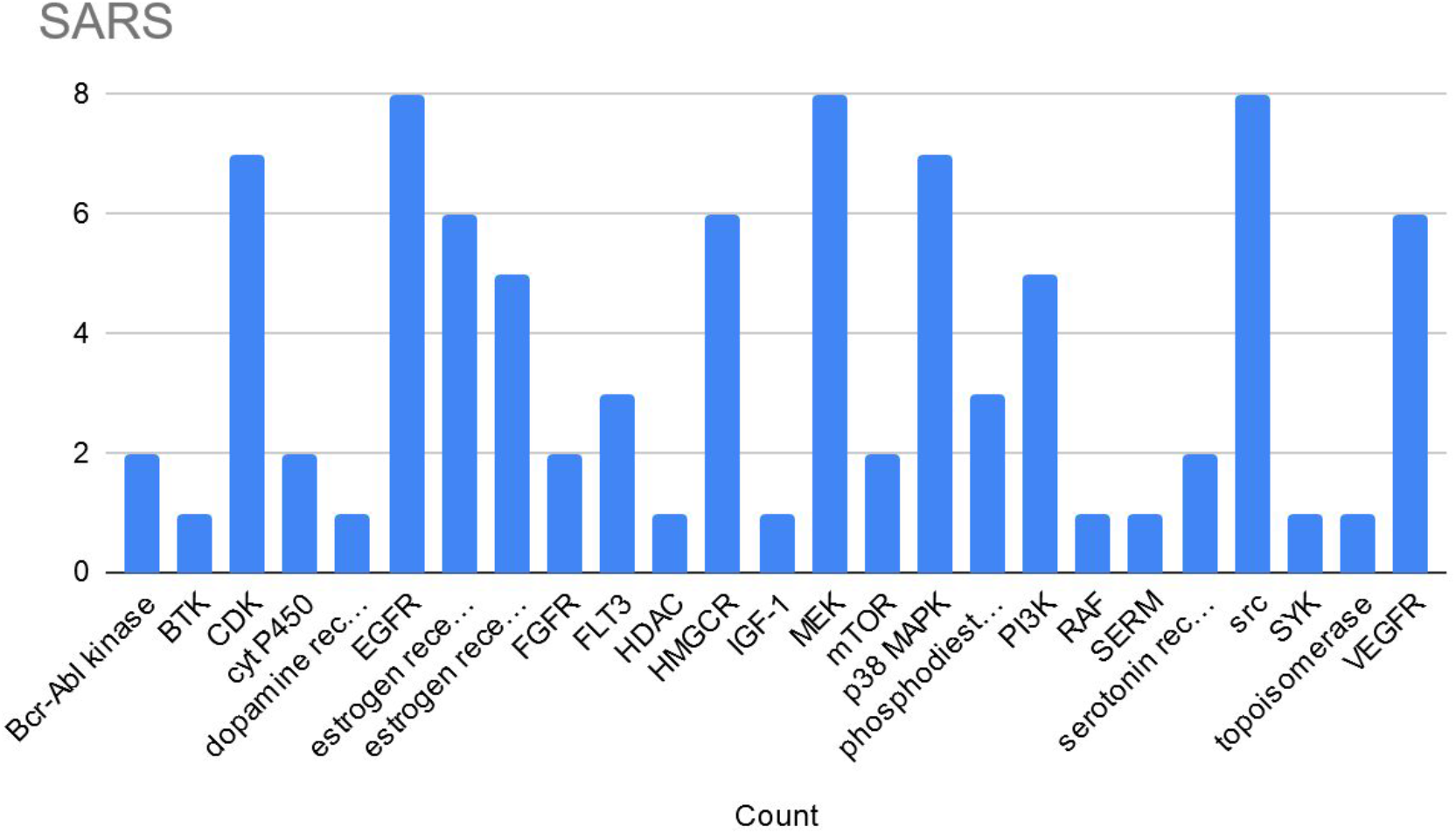
Anti-SARS-CoV-1 inhibitory mechanisms. Cumulative counts of the most significant potential inhibitory mechanisms against SARS-CoV-1 infections in nine separate samples.

SARS-CoV-2 infected samples displayed five mechanisms of action with inhibitory potential in at least three of the five samples analysed. These included CDK, EGFR, MEK and src inhibitors, as well as serotonin receptor antagonists, p38 MAPK inhibitors and 3-Hydroxy-3-Methylglutaryl-CoA Reductase (HMGCR) inhibitors (Fig 15).

**Fig 15.**
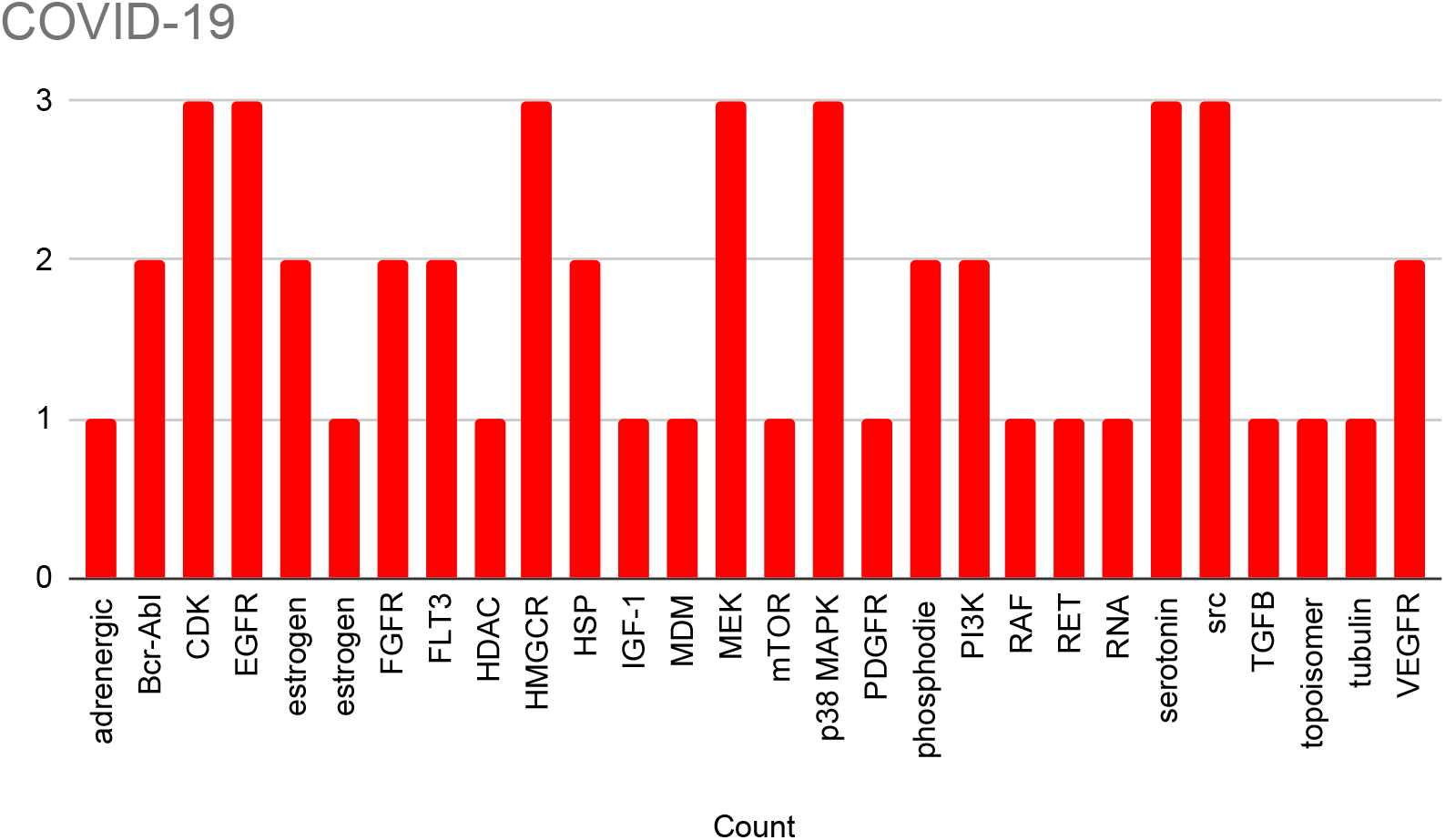
Anti-SARS-CoV-2 inhibitory mechanisms. Cumulative counts of the most significant potential inhibitory mechanisms against SARS-CoV-2 infections in five separate samples.

Overall, strong support for four mechanisms of action with broad anti-coronavirus inhibitory potential could be detected, namely: CDK, EGFR, MEK and src inhibitors.

Additionally, p38 MAPK inhibitors were also highly frequently negatively correlated with the gene expression profiles generated by SARS-CoV-1 and SARS-CoV-2 infections, while also being identified as potential inhibitors in three out of five MERS-CoV infected samples.

## Discussion

### Gene expression results

The number of differentially expressed genes varied greatly between samples, even when those were restricted only to samples collected at least 24h after infection. Indeed, the range could vary from 0 to almost 9000 genes. Even when keeping time point, species and host cell lines constant, as in the case of datasets GSE148729 and GSE37827, the number of differentially expressed genes ranged from 72 to 1835. Despite these differences, when clustering studies either by viral species or by host cell line, some macrotrends could be observed at the individual gene level.

An analysis of the expression of host genes or proteins involved in viral invasion (namely DPP4, ACE2 and TMPRSS2) provided a rather mixed picture. DPP4 appeared to be upregulated in Calu-3 cells, but downregulated in human fibroblasts (Fig 1A-3A). As a modulation of DPP4 by MERS viruses has not been implicated in any phenotypes, the actual biological significance of these observations or whether they represent a stochastic effect due to study setups, remains to be determined.

Conversely, ACE2 downregulation by the virus has been implicated in the pathology of SARS-CoV-1 [8] and potentially SARS-CoV-2 [34]. As expected, no alteration in expression was observed in MERS-infected cells. However, in SARS-CoV-1 and SARS-CoV-2 infected cells, a statistically significant downregulation of the gene could be observed only in one study and one cell line (Calu-3). Furthermore, in another study where Calu-3 cells were infected with SARS-CoV-1 (GSE37827), no significant downregulation was detectable (Fig 4B). There was no evidence of downregulation at the protein level either. Again, the discrepancies could be due to the many variables that separate each dataset. But it is also possible that ACE2 downregulation may only occur in certain cells *in vivo*, and requires an interaction with the host immune system *in vivo*. Alternatively, it could also be that the mouse model used to generate the original observation may not be a reliable model for human infections.

TMPRSS2 gene expression also provided a very mixed picture, with MERS studies showing both upregulation, downregulation and no difference, while in studies involving SARS-CoV-1 and SARS-CoV-2, a downregulation could be observed in Calu-3 cells only. Again, this may be more a reflection of the differences between studies than any biological significance.

Several cytokines have been implicated in the pathology caused by coronavirus infections. 15 of these were selected based on evidence in the literature and analysed for changes in expression at the gene level following viral infection. In the 6 transcriptomic studies, we found evidence for the significant upregulation of seven of these cytokines (namely IL-1B, IL-6, IL-8, IL-12A, CXCL10, CCL2 and TNF-alpha.) in at least a sample group per species. This is consistent with the theory of a “cytokine storm” occurring during coronavirus infections. On the other hand, there was a general lack of induction of type I interferons in SARS-CoV-1 and SARS-CoV-2 infections, particularly early on in the infection, consistent with observations that both viruses can interfere with the expression of these antiviral cytokines.

### Altered Pathway Analysis

Analysis of the proteome of cells infected with SARS-CoV-2 confirmed that the cholesterol pathway was downregulated, whereas carbon metabolism was upregulated, as described in the study by Bojkova et al [25]. No significant upregulation of the spliceosome could be observed, although this was most likely due to the more stringent analysis performed by the platform compared to traditional approaches for proteome analysis.

When looking at the transcriptome samples collected 24h post-infection, irrespective of cell-line, the upregulation of two KEGG pathways associated with immune responses was observed in all three viruses, as well as the upregulation of the “Ribosome biogenesis” pathway. The former two are expected as the results of defensive mechanisms against viral infections.The latter could be a consequence of the postulated disruption of ribosome biogenesis by the viral N protein [35]. The “propanoate metabolism” and “pyruvate metabolism” pathways were both downregulated in all three viruses and as they are both involved in carbohydrate metabolism, were in contrast with the findings from the proteome analysis. Conversely, the inhibition of the “peroxisome pathway” might reflect a direct exploitation by coronaviruses [36]. As the peroxisome also plays a role in cholesterol biosynthesis [37], this observation is consistent with the proteome data. When focusing only on the two SARS-CoV species, another pathway related to immune responses (“complement and coagulation cascades”) was upregulated in both species, while pathways related to carbohydrate metabolism were again downregulated. Additionally amino acid synthesis and drug metabolic pathways were also under-expressed in both viral species. Intriguingly, the disruption of the cytochrome P450 metabolic machinery in COVID-19 patients has been previously hypothesised [38].

Focusing the KEGG pathway analysis on samples with infected Calu-3 cells highlighted once more the upregulation of the “cytokine cytokine receptor” pathway in all three viral species. Several pathways were also downregulated in all three species. The downregulation of the “Lysosome” pathway may reflect the dependence of coronavirus cell entry on the lysosomal pathway [39]. Downregulation of the pyruvate pathway was related to the general inhibition of carbohydrate metabolism observed in the 24h samples. Finally, the downregulation of glutathione metabolism could reflect a direct viral manipulation of host defense mechanisms. Influenza infections have been associated with depletion of glutathione levels, which may favor viral replication [40]. The largest number of shared dysregulated pathways was observed between SARS-CoV-1 and SARS-CoV-2 infected Calu-3 cells. Besides immunity and cell stress related terms, upregulated pathways also highlighted the similarity with other viral diseases (Table 6). Pathways related to carbohydrate metabolisms were once more downregulated, highlighting the contrast between proteomic and transcriptomic datasets. The downregulation of the peroxisome pathway is consistent with glutathione inhibition, due to the role of the organelle in both the metabolism of reactive oxygen species and its antiviral properties [41]. The inhibition of the “butanoate metabolism” pathway could also fit this general trend of viral-mediated modulation of the immune system, due to its role in immune homeostasis [42].

### Drug Connectivity Map Analysis

Four main mechanisms of action (CDK, EGFR, MEK and src inhibitors) with broad inhibitory potential against human coronavirus infections were postulated across most of the datasets. An additional mechanism (p38 MAPK inhibitors) was also suggested when focusing on SARS-CoV-1 and SARS-CoV-2 infections. All these inhibitory mechanisms target kinases and indeed kinase inhibitors have been proposed as broad-spectrum antiviral therapies [43]. When looking at individual drugs that were identified as having anti-coronavirus properties by the bioinformatic platform, the MEK-inhibitor trametinib had already shown inhibitory potential against MERS-CoV infections [44]. Furthermore, various key cellular signaling pathways critical for SARS-CoV-2 infection have been identified [45]. These included the mTOR-PI3K-AKT and ABL-BCR/MAPK pathways, inhibitors against which also appeared in our analysis (Fig 12).

## Conclusion

Comparing gene expression coronavirus infection datasets across *in vitro* studies is fraught with potential pitfalls, due to the many variables that can alter the outcome of a study (such as viral strains used, infection protocols, cell lines used, time points of collection, sequencing technologies,etc..). Nonetheless, such information can provide some general insights on shared coronavirus infection features that can help discern potential targets for intervention and provide a better understanding of viral biology.

In this article, we observed several patterns shared across coronavirus species that were consistent with coronavirus biology and pathology. However, we also observed contrasting results (e.g. the expression of the ACE2 gene or regulation of the carbohydrate metabolism) that highlight the difficulty of working with heterogeneous datasets.

We could also identify shared potential mechanisms of viral inhibition that reflect what is known from the literature and further strengthen the argument for the selection of drugs that can be repurposed to treat coronavirus infections.

As a final note, all the datasets and analysis tools are currently available online (https://public.bigomics.ch/app/omicsplayground_viromics) and can be accessed to repeat and verify the analysis performed in this article.

## Supporting information

**S1 Table Dysregulated KEGG pathways across studies 24h post infection with MERS-CoV**. Combined significantly (q<0.05, logFC >0.5) dysregulated KEGG pathways across studies 24h post infection with MERS-Cov.

**S2 Table Dysregulated KEGG pathways across studies 24h post infection with SARS-CoV-1** Combined significantly (q<0.05, logFC >0.5) dysregulated KEGG pathways across studies 24h post infection with SARS-Cov-1.

**S3 Table Dysregulated KEGG pathways across studies 24h post infection with SARS-CoV-2** Combined significantly (q<0.05, logFC >0.5) dysregulated KEGG pathways across studies 24h post infection with SARS-Cov-2.

**S4 Table Dysregulated KEGG pathways in Calu-3 cells infected with MERS-CoV.** Combined significantly (q<0.05, logFC >0.5) dysregulated KEGG pathways across studies in Calu-3 cells infected with MERS-Cov.

**S5 Table Dysregulated KEGG pathways in Calu-3 cells infected with SARS-CoV-1.** Combined significantly (q<0.05, logFC >0.5) dysregulated KEGG pathways across studies in Calu-3 cells infected with SARS-Cov-1.

**S6 Table Dysregulated KEGG pathways in Calu-3 cells infected with SARS-CoV-2.** Combined significantly (q<0.05, logFC >0.5) dysregulated KEGG pathways across studies in Calu-3 cells infected with SARS-Cov-2.

## Notes

### Competing Interest Statement

The authors have declared no competing interest.

